# TRIM9-dependent ubiquitination of DCC constrains kinase signaling, exocytosis, and axon branching

**DOI:** 10.1101/154666

**Authors:** Melissa Plooster, Shalini Menon, Cortney C. Winkle, Fabio L. Urbina, Caroline Monkiewicz, Kristen D. Phend, Richard J. Weinberg, Stephanie L. Gupton

**Author notes:** contributed equally to this work (Melissa Plooster, Shalini Menon). 111 Mason Farm Road, CB 7090, Chapel Hill NC 27599.

## Abstract

Extracellular netrin-1 and its receptor DCC promote axon branching in developing cortical neurons. Netrin-dependent morphogenesis is preceded by multimerization of DCC, activation of FAK and Src family kinases, and increases in exocytic vesicle fusion, yet how these occurrences are linked is unknown. Here we demonstrate that TRIM9-dependent ubiquitination of DCC blocks the interaction with and phosphorylation of FAK. Upon netrin-1 stimulation TRIM9 promotes DCC multimerization, but TRIM9-dependent ubiquitination of DCC is reduced, which promotes an interaction with FAK and subsequent FAK activation. We found that inhibition of FAK activity blocks elevated frequencies of exocytosis *in vitro* and elevated axon branching *in vitro* and *in vivo.* Although FAK inhibition decreased SNARE-mediated exocytosis, assembled SNARE complexes and vesicles adjacent to the plasma membrane were increased, suggesting a novel role for FAK in the progression from assembled SNARE complexes to vesicle fusion in developing murine neurons.

**Abbreviations used in this paper:**

DCC: Deleted in Colorectal Cancer
TRIM: Tripartite Motif
SFK: src family kinase
DCC^KR^: non ubiquitinatable DCC mutant
VAMP: vesicle associated membrane protein
TRIM9ΔRING: TRIM9 lacking the ubiquitin ligase RING domain
TRIM9ΔSPRY: TRIM9 variant lacking the DCC-binding SPRY domain
TIRF: Total Internal Reflection Fluorescence
pY: phosphotyrosine
FAKi: pharmacological FAK inhibitor 14
FRNK: FAK related non-kinase
STX-1A: syntaxin 1A
IP: immunoprecipitate

## Introduction

During development, extracellular axon guidance cues direct the extension and branching of axons, essential for appropriate anatomy and function in the adult brain. Disruption of axon extension and branching may lead to the defective connectivity implicated in neurodevelopmental and neuropsychiatric disorders (Engle, 2010; Grant *et al.*, 2012). In the mammalian neocortex, the secreted axon guidance cue netrin-1 promotes attractive guidance and axon branching through the receptor Deleted in Colorectal Cancer (DCC) (Kennedy and Tessier-Lavigne, 1995; Keino-Masu *et al.*, 1996). Gene trap-mediated disruption or deletion of either *Dcc* or the gene encoding netrin-1 (*Ntn1*) in mice leads to defects in fiber tracts in the forebrain, including the corpus callosum and hippocampal commissure, and perinatal or embryonic lethality (Serafini *et al.*, 1996; Fazeli *et al.*, 1997; Bin *et al.*, 2015; Yung *et al.*, 2015). Netrin-dependent neuronal morphogenesis is preceded by several subcellular events. In the presence of netrin-1, DCC is recruited to clusters within the plasma membrane where it homomultimerizes (Mille *et al.*, 2009; Matsumoto and Nagashima, 2010; Wang *et al.*, 2014; Gopal *et al.*, 2016). The noncatalytic cytoplasmic tail of DCC interacts with non-receptor tyrosine kinases, leading to their activation. These targets include FAK and Src family kinases (SFKs), which are involved in cell adhesion, migration survival, neuritogenesis, and axon outgrowth (Li *et al.*, 2004; Liu *et al.*, 2004; Meriane *et al.*, 2004; Ren *et al.*, 2004). The molecular regulators that control and coordinate DCC relocalization and downstream kinase activation are not fully elucidated.

We identified vertebrate TRIM9, an evolutionarily conserved class I TRIpartite Motif (TRIM) protein (Berti *et al.*, 2002; Tanji *et al.*, 2010), as a key regulator of netrin-dependent morphogenesis in cortical and hippocampal neurons (Winkle *et al.*, 2014; Menon and Boyer *et al.*, 2015; Winkle and Olsen *et al.*, 2016). The single invertebrate orthologs of *Ntn1* and of *Dcc* regulate axon development through the single class I TRIM ortholog (Hao *et al.*, 2010; Morikawa *et al.*, 2011). Mammalian TRIM9 directly interacts with the cytoplasmic tail of DCC, the neuronal exocytic t-SNARE SNAP25, and the filopodial actin polymerase VASP (Li *et al.*, 2001; Winkle *et al.*, 2014; Menon and Boyer *et al.*, 2015). Genetic deletion of murine *Trim9* in cortical neurons is associated with a loss of netrin-1 responsiveness, elevated exocytosis, enhanced growth cone filopodial stability *in vitro*, and defects in axon branching and axon projections *in vitro* and *in vivo* (Winkle *et al.*, 2014; Menon *et al.*, 2015; Winkle and Olsen *et al.*, 2016). However, whether TRIM9 plays a role in DCC localization or FAK-dependent intracellular signal transduction downstream of DCC/netrin-1 remains unknown.

Here we find that DCC is ubiquitinated in a TRIM9-dependent manner. Our data are consistent with the hypothesis that DCC ubiquitination blocks the activation of FAK and SFK in the absence of netrin-1. Following netrin-1 stimulation, DCC ubiquitination is reduced, and TRIM9-dependent clustering and multimerization of DCC occurs. FAK also becomes phosphorylated and activated, and SNARE-mediated exocytosis and axon branching increase. Inhibition of FAK activity blocks netrin-dependent exocytosis and axon branching but surprisingly, increases the amount of assembled SNARE complexes and the density of vesicles found immediately adjacent to the plasma membrane. This suggests a novel requirement for FAK activity in the progression from an assembled SNARE complex to SNARE-mediated fusion, which is necessary for plasma membrane expansion during axon branching.

## Results

### Deletion of *Trim9* disrupts netrin-dependent clustering and multimerization of DCC

Application of netrin-1 induces clustering of its receptor DCC at the plasma membrane (Matsumoto and Nagashima, 2010; Gopal *et al.*, 2016). Since we had observed colocalization of TRIM9 and DCC in punctae along the periphery of neurites (Winkle *et al.*, 2014), we hypothesized that TRIM9 may play a role in netrin-dependent DCC clustering. Using total internal reflection fluorescence (TIRF) microscopy, we observed rapid mCherry-DCC cluster formation at the cell edge of *Trim9^+/+^* neurons immediately following netrin-1 stimulation, which was absent in *Trim9^-/-^* neurons (**Fig. 1A, Movie 1**). To quantify the relocalization of DCC, we compared the change in mCherry-DCC fluorescence at the cell edge (**Fig1B,** outer 0.64 μm, blue) and the cell center (green), normalized to the whole cell (**Fig. 1B**, red). To reveal trends in fluorescence intensity changes, these data were fit to a local polynomial regression (**Fig. 1C**). In *Trim9^+/+^* neurons, DCC fluorescence increased at the cell edge and decreased in the cell center after netrin-1 stimulation, suggesting a rapid rearrangement of DCC. Clustered mCherry-DCC should exhibit higher fluorescence intensity than non-clustered DCC. To quantify changes in the localization of clustered DCC, we considered the localization of the brightest 10% of mCherry-DCC containing pixels (**Fig. 1D**). This revealed that the brightest DCC containing pixels (likely DCC clusters) increasingly localized to the cell edge following netrin-1 stimulation in *Trim9^+/+^* neurons. Netrin-dependent clustering of DCC was also observed on the apical plasma membrane by spinning disk confocal microscopy (**Fig. S1)**, indicating that DCC relocalization was not limited to the basal cell surface.

**Figure 1.**
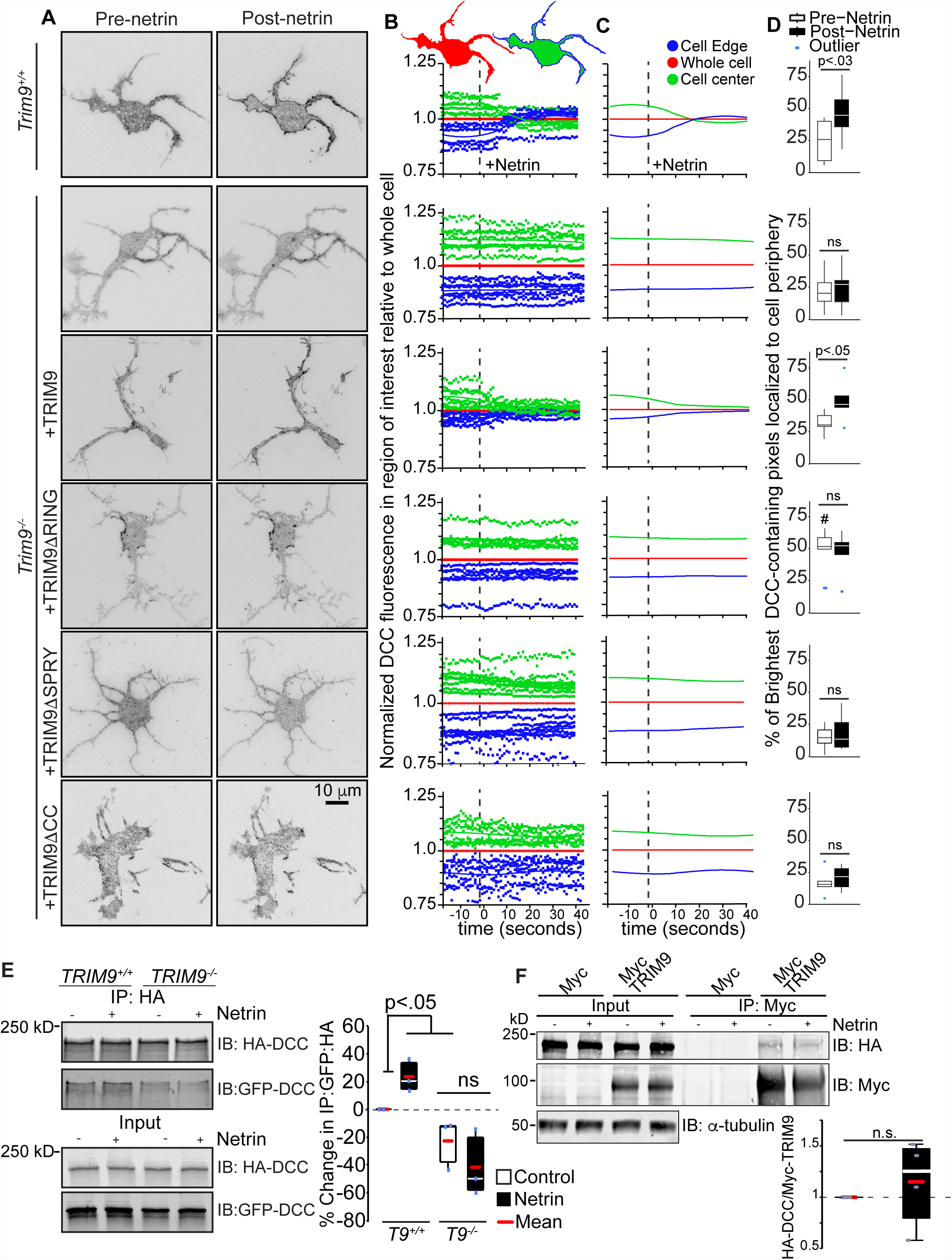
Loss of *Trim9* impairs netrin-dependent clustering and multimerization of DCC. **A)** Inverted TIRF images of *Trim9^+/+^* and *Trim9^-/-^* cortical neurons expressing fluorescently-tagged DCC before and after netrin-1 application. **B)** Schematic of cell mask showing whole cell (red), cell edge (blue, 0.64 μm) and cell center (green) with graphs of average mean fluorescence of DCC in the denoted ROIs in individual cells over time. Values are normalized to the average intensity of the whole cell. **C)** Local polynomial regression of data shown in B) demonstrates trends in temporal changes in average DCC fluorescence intensity in each region of interest. **D)** The percent of the pixels containing the highest 10% of DCC fluorescence intensity in the TIRF illumination field that localize at the cell perimeter. p values assessed via two way ANOVA with Tukey post-hoc correction show differences between each pre and post netrin-1 condition. # denotes p<.0005 difference from pre-netrin-1 *Trim9^+/+^*. **E)** Immunoblot of inputs and immunoprecipitates (IP) of HA-DCC. Co-IP of GFP-DCC increases following netrin-1 stimulation in *TRIM9^+/+^* cells. p values assessed by Kruskal-Wallis ANOVA with LSD post-hoc correction. **F)** Immunoblot of inputs and immunoprecipitates (IP) of Myc and Myc-TRIM9 in *TRIM9^+/+^* HEK 293 cells. p values assessed by Mann-Whitney test.

This netrin-dependent redistribution and clustering of mCherry-DCC was absent in *Trim9^-/-^* neurons (**Fig. 1A-D, Movie 1**) and rescued by reintroduction of eGFP-TRIM9 (**Fig. 1A-D, Movie 2,** arrowheads), indicating that DCC clustering was dependent upon TRIM9. Expression of a TRIM9 mutant lacking the ubiquitin ligase RING domain (TRIM9ΔRING) induced clustering of DCC independent of netrin-1 (**Fig. 1A-D, Movie 2**). This gain-of-function suggests that the ligase domain of TRIM9 inhibits clustering of DCC in the absence of netrin-1. Expression of a TRIM9 variant lacking the DCC-binding SPRY domain (TRIM9ΔSPRY), or a mutant lacking the TRIM multimerization coiled-coil motif (TRIM9ΔCC), failed to rescue netrin-dependent clustering of DCC (**Fig. 1A-D, Movie 2**), suggesting that the interaction between TRIM9 and DCC and the multimerization of TRIM9 are both critical for TRIM9-dependent DCC clustering. These results reveal a netrin-stimulated, TRIM9-dependent spatial reorganization of DCC at the plasma membrane.

In addition to clustering in the plasma membrane, DCC homo-multimerizes upon application of netrin-1, as revealed by the co-immunoprecipitation of differentially-tagged DCC proteins (Mille *et al.*, 2009; Bin *et al.*, 2013). Since spatial clustering may involve multimerization (**Fig. 1A-D)**, we hypothesized that TRIM9 may also be involved in DCC multimerization. Using previously reported *TRIM9^+/+^* and *TRIM9^-/-^* HEK293 cells (Menon and Boyer *et al.*, 2015), we found that immunoprecipitation of HA-DCC with antibodies against the HA epitope co-precipitated pHluorin-DCC (**Fig. 1E**). Co-immunoprecipitation of pHluorin-DCC was enhanced by netrin-1 stimulation in *TRIM9^+/+^* HEK293 cells and was reduced and netrin-insensitive in *TRIM9^-/-^* HEK293 cells. Thus multimerization of DCC was promoted by TRIM9. We did not observe netrin-dependent changes in the amount of DCC co-immunoprecipitated by TRIM9 (**Fig. 1F**), indicating that netrin-dependent changes in DCC multimerization cannot be explained by an increased interaction between TRIM9 and DCC.

### Loss of *Trim9* leads to hyperactivation of FAK

We hypothesized that the defects in netrin-dependent clustering and multimerization of DCC may be associated with defects in the netrin-triggered activation of FAK and SFK. Previous studies have shown that the non-receptor tyrosine kinase FAK directly interacts with the COOH-terminal LD motif on the P3 domain of DCC (**Fig. 2A**) and that FAK is autophosphorylated at Y397 (pY397) upon netrin stimulation, leading to FAK activation (Li *et al.*, 2004; Liu *et al.*, 2004; Ren *et al.*, 2004). We similarly observed netrin-dependent increases in FAK pY397 in *Trim9^+/+^* neurons and, surprisingly, found enhanced basal FAK pY397 in *Trim9^-/-^* neurons (**Fig. 2B)**. Although netrin-dependent clustering of DCC is associated with activation of FAK, these data suggest clustering is not necessary for FAK activation and further, that TRIM9 constrains FAK activation in the absence of netrin.

**Figure 2.**
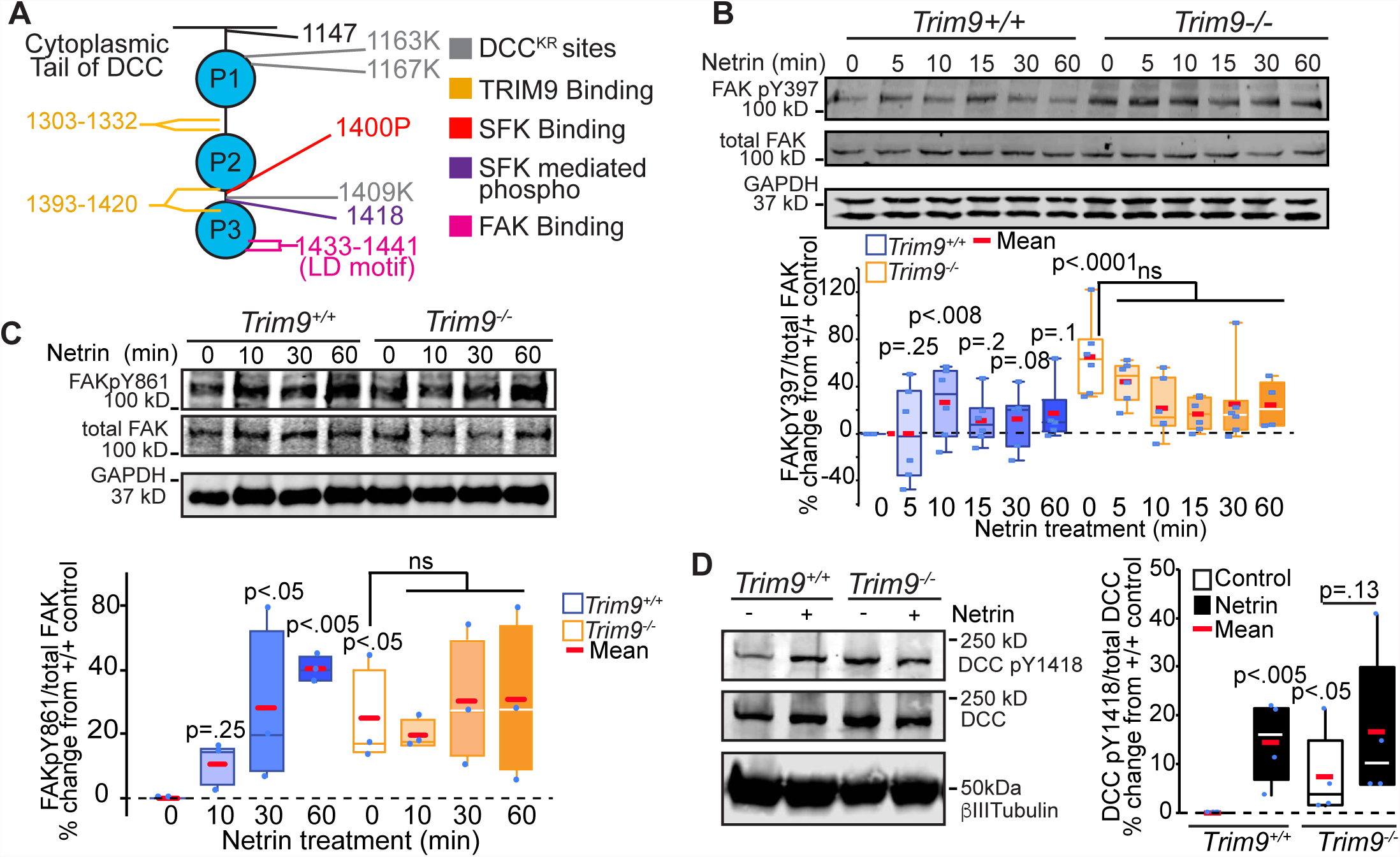
Loss of *Trim9* leads to hyperactivation of FAK. **A)** Schematic representation of the cytoplasmic tail of DCC showing key mutation and interaction sites. **B)** Immunoblots of endogenous total FAK, FAKpY397 and GAPDH from cortical lysate. Quantification of percent change in each phosphorylation event normalized to total FAK relative to *Trim9^+/+^* neurons. p values noted for Kruskal-Wallis ANOVA with LSD post-hoc correction to *Trim9^+/+^* control, unless otherwise noted. **C)** Immunoblots of endogenous total FAK, FAKpY861 and GAPDH from cortical lysate. Quantification of percent change in each phosphorylation event normalized to total FAK relative to *Trim9^+/+^* neurons. p values noted for Kruskal-Wallis ANOVA with LSD post-hoc correction to *Trim9^+/+^* control, unless otherwise noted. **D)** Immunoblots of endogenous DCC pY1418, DCC and βIII tubulin in cortical neurons. Quantification of levels of pY1418/DCC relative to *Trim9^+/+^* neurons. p values noted for Kruskal-Wallis ANOVA with LSD post-hoc correction compared to *Trim9^+/+^* control, unless otherwise noted.

SFKs dock at pY397 on FAK and interact with the PXXP motif beginning at P1400 on the cytoplasmic tail of DCC (Li *et al.*, 2004; Ren *et al.*, 2004, **Fig. 2A**). This interaction leads to a number of SFK-dependent phosphorylation events, including FAK pY861 and DCC pY1418. In *Trim9^+/+^* neurons, these SFK-dependent phosphorylation events increased in response to netrin-1 stimulation (**Fig. 2C-D**), recapitulating previous results (Li *et al.*, 2004; Liu *et al.*, 2004; Meriane *et al.*, 2004). However, these phosphorylation events were aberrantly elevated in *Trim9^-/-^* cortical neurons, suggesting that FAK hyperactivation promotes downstream phosphorylation events. TRIM9 interacts with two sites on the cytoplasmic tail of DCC (Winkle *et al.*, 2014). These sites do not overlap with the established FAK binding site (**Fig. 2A**), and we did not observe reduced interaction between TRIM9 and DCC upon netrin addition (**Fig. 1F**), suggesting that a different TRIM9-dependent mechanism controls activation of FAK.

### TRIM9-dependent ubiquitination of DCC does not alter protein stability

Because TRIM9 is a brain-enriched E3 ubiquitin ligase, and ubiquitination of DCC in embryonic cortical neurons and heterologous cells has been reported (Hu *et al.*, 1997; Kim *et al.*, 2005), we examined whether DCC ubiquitination was dependent on TRIM9. In *TRIM9^+/+^* HEK293 cells, immunoprecipitation of HA-DCC under denaturing conditions revealed a HA+ band at the expected molecular weight (~190, **Fig. 3A,** red arrowhead) and a heavier HA+ band that migrated with co-immunoprecipitated FLAG-ubiquitin (**Fig. 3A,** blue arrowhead). These higher molecular weight species are indicative of DCC ubiquitination. Either netrin-1 addition in the presence of TRIM9 or genetic loss of *TRIM9* reduced FLAG-ubiquitin co-immunoprecipitated by HA-DCC (**Fig. 3A)**, suggesting that this modification is netrin-sensitive and TRIM9-dependent. To confirm the relevance of these results in neurons, we immunoprecipitated endogenous DCC from primary cultured *Trim9*^+/+^ or *Trim9*^-*/-*^ embryonic cortical neurons under denaturing conditions and immunoblotted for DCC and ubiquitin (**Fig. 3B)**. This revealed that cortical neurons also exhibited TRIM9-dependent, netrin-sensitive DCC ubiquitination. SDSPAGE and immunoblotting of cortical neuron lysate resolved a DCC immunoreactive band at the expected molecular weight of 190 kD (**Fig. 3C**). However, neither netrin addition, genetic loss of *Trim9,* nor treatment with the proteosome inhibitor Bortezomib altered the level of this band, suggesting that TRIM9-dependent ubiquitination of DCC does not alter DCC stability.

**Figure 3.**
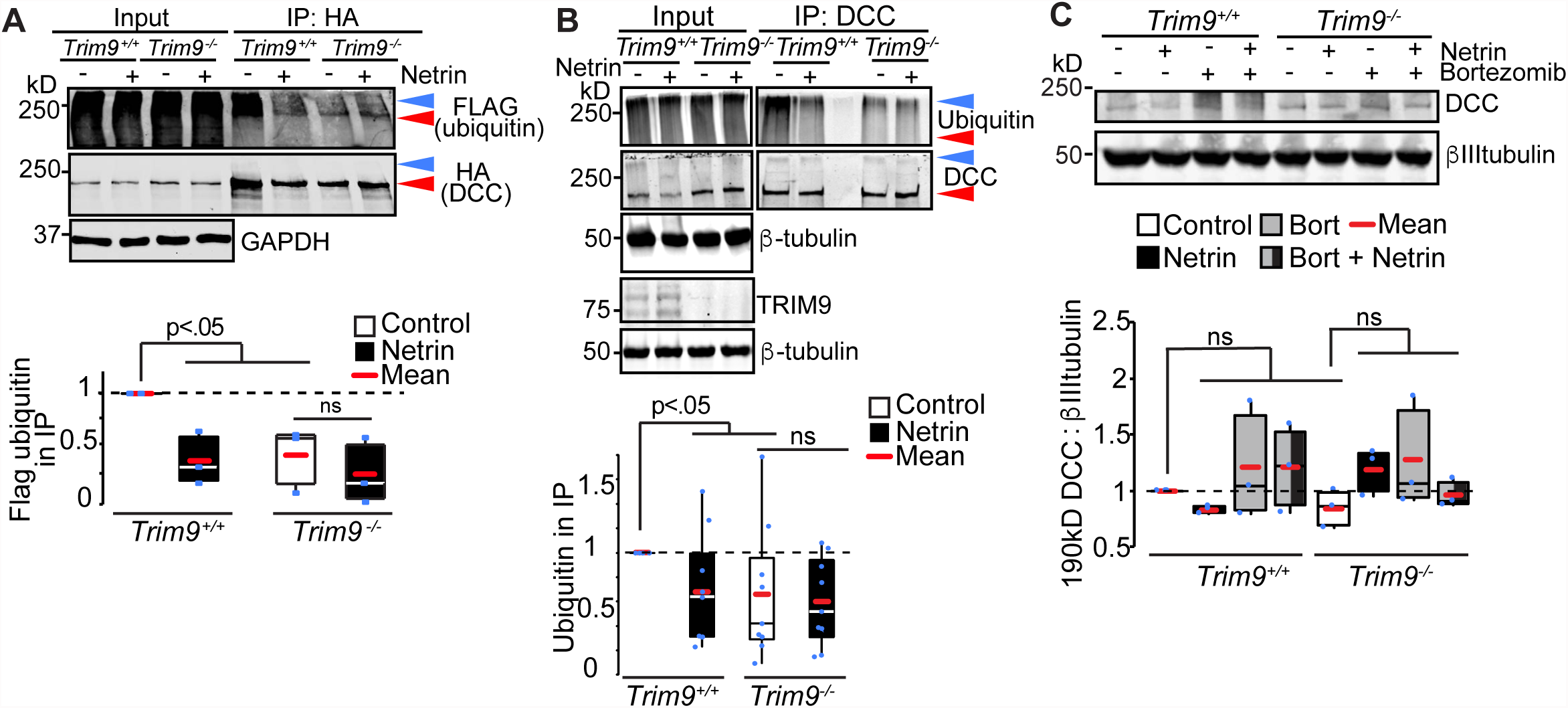
Netrin-sensitive, TRIM9-dependent ubiquitination of DCC. **A)** SDS-PAGE and western blots of inputs and HA-DCC immunoprecipitates (IP) from *TRIM9^+/+^* and *TRIM9^-/-^* HEK293 cells blotted for Flag-ubiquitin and HA-DCC. High molecular weight ubiquitin in the IP is considered DCC ubiquitination (blue arrowhead). Red arrowhead demarcates DCC at expected molecular weight ~190. p values from Kruskal-Wallis ANOVA with LSD post-hoc correction. **B)** SDS-PAGE and western blots of inputs and DCC immunoprecipitates (IP) from *Trim9^+/+^* and *Trim9^-/-^* cortical neurons blotted for endogenous ubiquitin and DCC. Blue arrowhead demarcates ubiquitinated DCC, red arrowhead demarcates DCC at expected molecular weight ~190. p values from Kruskal-Wallis ANOVA with LSD post-hoc correction compared to *Trim9^+/+^* control. **C)** Immunoblot of cortical lysate shows DCC at expected molecular weight (~190 kD) is insensitive to inhibition of the proteasome (100 μM bortezomib) or deletion of *Trim9.*

### A DCC mutant lacking ubiquitination sites shows increased interaction with FAK and increased phosphotyrosine levels

Bioinformatic analysis of the cytoplasmic tail of DCC identified three residues (K1163, K1167, and K1409) as putative ubiquitination sites, which we mutated to R (DCC^KR^). One of these K residues (1409K) is situated between the FAK and SFK binding sites, near Y1418 on DCC (**Fig. 2A**). pHluorin-DCC^KR^ exhibited reduced ubiquitination in HEK293 cells compared to pHluorin-DCC (**Fig. 4A**). Based on the size of ubiquitin (~8 kD) relative to the P3 domain of DCC (~3.7 kD) and our observation that FAK is activated upon reduced DCC ubiquitination (**Fig. 2B, 3A-B**), we hypothesized that ubiquitination of DCC occludes access of FAK to the P3 domain and prevents the downstream activation of FAK and SFK. Consistent with this hypothesis, compared to pHluorin-DCC, pHluorin-DCC^KR^ exhibited increased interaction with Myc-FAK in HEK293 cells, as measured by co-immunoprecipitation (**Fig. 4B)**. Further in *TRIM9^+/+^* HEK293 cells, DCC pY1418 was enhanced on pHluorin-DCC^KR^ as compared to pHlourin-DCC (**Fig. 4C**). In *TRIM9^-/-^* cells pY1418 DCC was enhanced compared to *TRIM9^+/+^* HEK293 cells and insensitive to netrin-1 or the K-R mutations. These data suggest that TRIM9-dependent ubiquitination of DCC blocks access of FAK to DCC, which inhibits downstream FAK and SFK-dependent phosphorylation events. pHluorin-DCC^KR^ did display rapid netrin-dependent clustering at the periphery of *Trim9^+/+^* neurons (**Fig. 4D, Movie 3)**, indicating that netrin-dependent clustering is independent of changes in the status of DCC ubiquitination.

**Figure 4.**
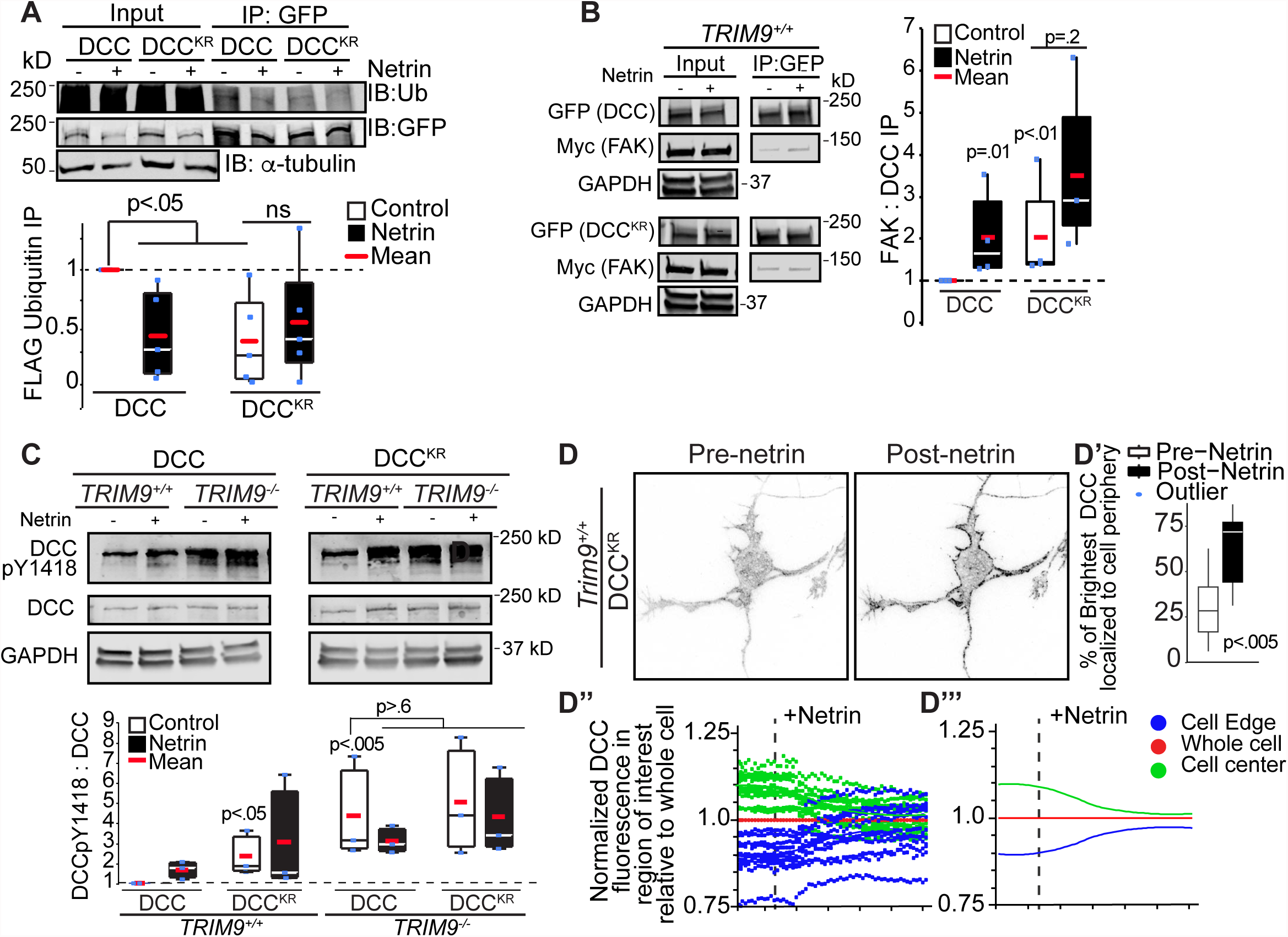
Ubiquitination of DCC regulates FAK interactions and activation, but not DCC clustering. **A)** SDS-PAGE and western blots of inputs and GFP-DCC immunoprecipitates (IP) from *TRIM9^+/+^* HEK293 cells blotted for Flag-ubiquitin and GFP-DCC. High molecular weight ubiquitin in the IP is considered DCC ubiquitination. p values from Kruskal-Wallis ANOVA with LSD post-hoc correction. **B)** Western blot of inputs and immunoprecipitates (IP) of GFP-DCC or GFP-DCC^KR^ and Myc-FAK from *TRIM9^+/+^* and *TRIM9^-/-^* HEK293 cells. p values noted for Kruskal-Wallis ANOVA with LSD post-hoc correction. **C)** Immunoblots of DCC pY1418, DCC and GAPDH in HEK293 cells expressing GFP-DCC or GFP-DCC^KR^. Basal levels of pY1418 are increased on DCC^KR^ and in *TRIM9^-/-^* cells. p values noted for Kruskal-Wallis ANOVA with LSD post-hoc compared to DCC in *TRIM9^+/+^* control. **D)** Inverted TIRF images of *Trim9^+/+^* cortical neuron expressing fluorescently-tagged DCC^KR^ before and after netrin-1 application. **D’)** The percent of the pixels containing the highest 10% of DCC fluorescence intensity in the TIRF illumination field that localize at the cell perimeter. p values assessed via t-test show differences between each pre-and post netrin-1 condition. **D”)** Graphs of average mean fluorescence of DCC in the denoted ROIs in individual cells over time. Values are normalized to the average intensity of the whole cell within the first 15 frames of imaging. **D”’)** Local polynomial regression of data demonstrates trends in temporal changes in average DCC fluorescence intensity in each ROI.

### Inhibition of FAK activity blocks axon branching

Netrin-1 promotes axon branching, whereas deletion of *Trim9* aberrantly elevates branching in a netrin-independent manner (Dent *et al.*, 2004; Winkle *et al.*, 2014). FAK and SFK activity are required for netrin-dependent axon outgrowth and guidance (Liu *et al.*, 2004; Ren *et al.*, 2004; Mehlen and Rama, 2007) but have not been implicated in axon branching. To determine whether FAK activity was required for netrin-dependent axon branching and the elevated axon branching observed in *Trim9^-/-^* neurons, we used a pharmacological inhibitor, FAK Inhibitor 14 (FAKi), which prevents autophosphorylation of FAK at Y397 (Golubovskaya *et al.*, 2008) and thus FAK activity. FAKi treatment blocked netrin-dependent increases in axon branching density in *Trim9^+/+^* neurons and reduced the exuberant axon branching phenotype in *Trim9^-/-^* neurons (**Fig. 5A**), without significant alterations to axon length. Similar results were observed upon expression of FAK-related non-kinase (FRNK, **Fig. 5B**), a noncatalytic variant of FAK that acts as a dominant negative (Richardson and Parsons, 1996). These data suggest that FAK activity is required downstream of DCC and TRIM9 to promote netrin-dependent axon branching and for the increased axon branching that occurs in the absence of *Trim9*.

**Figure 5.**
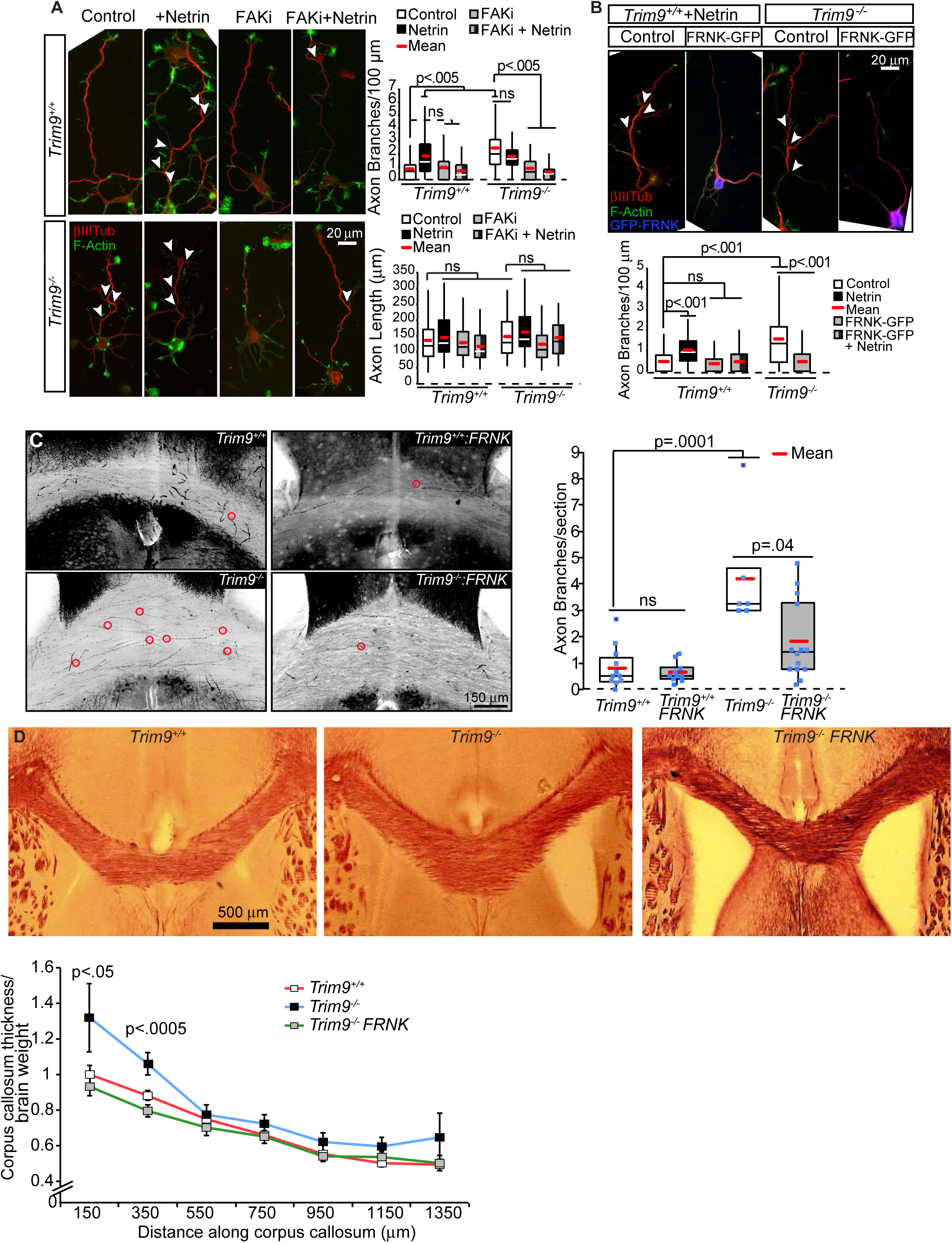
FAK activity is required for axon branching. **A)** Phalloidin-stained F-actin (green) and βIII tubulin (red) in *Trim9^+/+^* and *Trim9^-/-^* cortical neurons at 72 h in culture. Quantification of density of axon branches/ 100 μm axon length. p values from Kruskal-Wallis ANOVA with Bonferroni post-hoc correction. **B)** Phalloidin-stained F-actin (green), βIII tubulin (red) +/- GFP-FRNK (blue) in *Trim9^+/+^* and *Trim9^-/-^* cortical neurons 72 h after plating. Quantification of density of axon branches/100 μm axon length. p values from Kruskal-Wallis ANOVA with Bonferroni post-hoc correction. **C)** Inverted maximal projections of *Trim9^+/+^*/Thy1-GFP, *Trim9^+/+^*/Thy1-GFP/NexCre/FRNK^LoxSTOPLox^, *Trim9^-/-^*/Thy1-GFP and *Trim9^-/-^*/Thy1-GFP/NexCre/FRNK^LoxSTOPLox^ corpus callosum sections. Red circles denote branch points. Quantification of average axonal branches per section per mouse, n = 7-15 mice/genotype. p values from Kruskal-Wallis ANOVA with Bonferroni post-hoc correction. **D)** Black gold staining and quantification of position-matched coronal sections through *Trim9^+/+^*, *Trim9^-/-^*/Thy1-GFP and *Trim9^-/-^*/Thy1-GFP/NexCre/FRNK^LoxSTOPLox^ mice. Quantification of density of axon branches normalized to brain weight.p values from Kruskal-Wallis ANOVA.

Using a Thy1-GFP reporter mouse (Feng *et al.*, 2000), we previously found that *Trim9^-/-^* mice exhibit aberrant axon branching within the corpus callosum (Winkle *et al.*, 2014), which was associated with corpus callosum thickening. To test whether FAK activity was required for exuberant branching *in vivo,* we crossed *Trim9^-/-^*/Thy1-GFP mice with FRNK^LoxSTOPLox^ (DiMichele *et al.*, 2009) and Nex:CRE mice (Goebbels *et al.*, 2006). The resultant mice express Cre in postmitotic cortical neurons, which promotes recombination and expression of FRNK specifically in these cells. Z-stacks of position-matched coronal sections through the corpus callosum were imaged by laser scanning confocal microscopy, and GFP+ axons were inspected for branch points. FRNK expression did not affect axon branching in the corpus callosum of *Trim9^+/+^* mice but did reduce the aberrant axon branching in the corpus callosum of *Trim9^-/-^* mice (**Fig. 5C, Movie 4)**, suggesting that FAK activity is necessary for aberrant axon branching in the absence of *Trim9 in vivo*. Myelin staining of position-matched coronal sections revealed that FRNK expression also reduced the elevated thickness in the rostral portion of the *Trim9^-/-^* corpus callosum (**Fig. 5D)**, further supporting the hypothesis that this thickening is at least partially due to aberrant axon branching.

### FAK inhibition blocks SNARE-dependent exocytosis but not SNARE complex assembly

Axon branching increases the neuronal plasma membrane surface area, which necessitates insertion of new plasma membrane material (Winkle *et al.*, 2016b). We previously demonstrated that netrin-dependent axon branching and the exuberant axon branching in *Trim9^-/-^* neurons requires SNARE-mediated exocytosis and is associated with increased SNARE complex assembly and an increased frequency of exocytosis (Winkle *et al.*, 2014). Since FAK inhibition blocked axon branching (**Fig. 5**), we hypothesized that FAK activation promotes the requisite SNARE complex assembly and exocytosis, thus driving plasma membrane expansion during axon branching. We exploited FAKi treatment and the SDS-resistance of SNARE complexes (Söllner *et al.*, 1993; Hayashi *et al.*, 1995) to investigate this possibility (**Fig. 6A)**. Netrin-1 stimulation of *Trim9^+/+^* neurons or deletion of *Trim9* increased the amount of assembled SNARE complexes (**Fig. 6A**) as previously observed. Unexpectedly, FAKi treatment increased basal levels of assembled SNARE complexes in *Trim9^+/+^* neurons and failed to disrupt the elevated SNARE assembly in *Trim9^-/-^* neurons. These data suggested that FAK activity is not required for SNARE complex formation. To determine if the elevated SNARE complexes that were present following FAKi treatment were driving increased exocytosis, we tracked the frequency of VAMP2-phluorin vesicle fusion events via TIRF microscopy. Application of netrin-1 increased basal exocytic fusion frequency in *Trim9^+/+^* cortical neurons as expected, however this netrin-1 dependent response was blocked by FAKi treatment (**Fig. 6B, Movie 5)**. Similarly, the aberrantly high exocytic vesicle fusion in *Trim9^-/-^* neurons was decreased by FAKi treatment (**Fig. 6B**).

**Figure 6.**
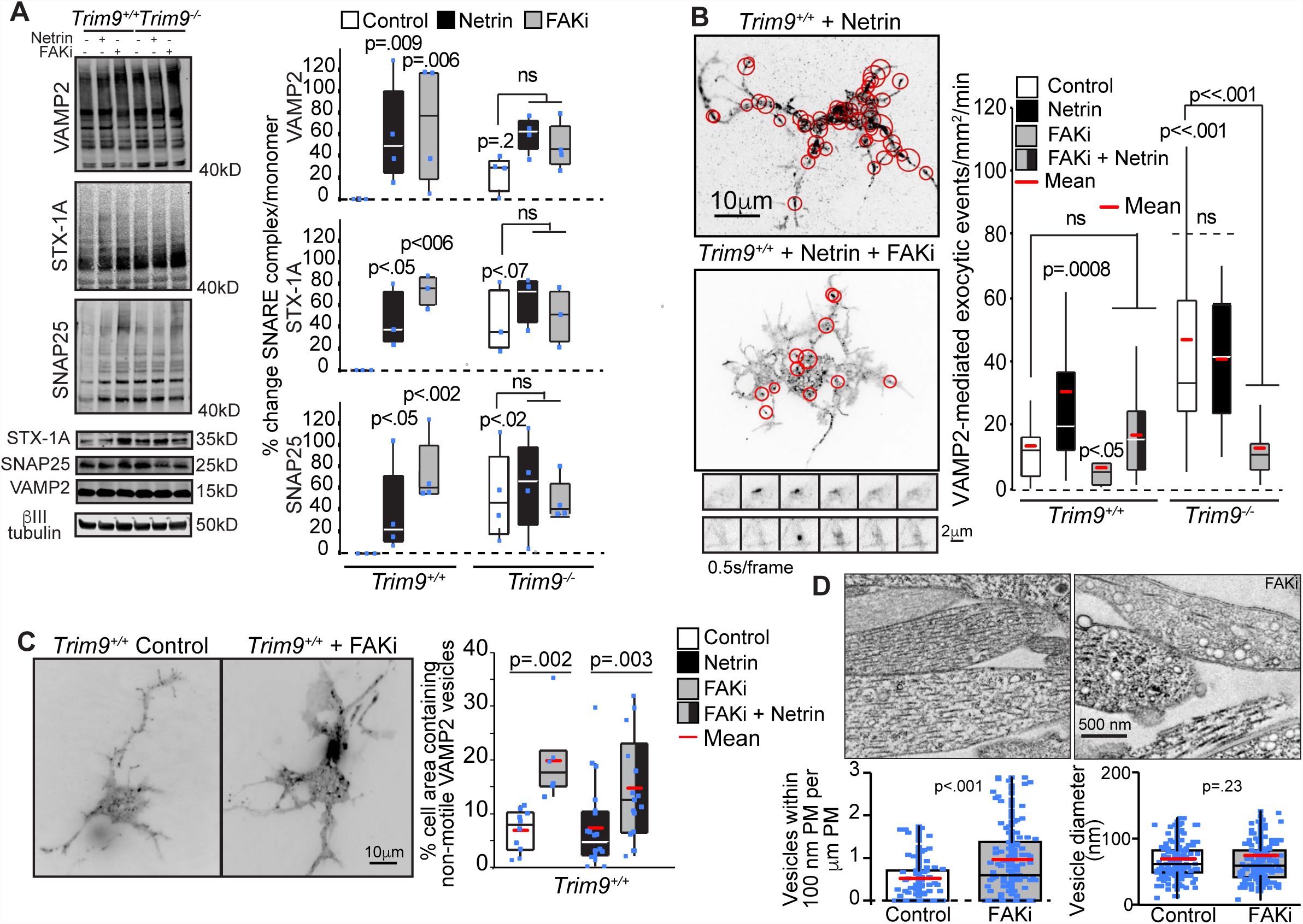
Inhibition of FAK activity blocks SNARE-mediated exocytic vesicle fusion but not SNARE complex formation. **A)** Immunoblots and quantification of SNARE complexes (above 40 kD), and monomers for VAMP2, Syntaxin-1 (STX-1A), and SNAP25. p values from Kruskal-Wallis ANOVA with LSD post-hoc correction compared to *Trim9^+/+^* control, unless otherwise noted. **B)** Quantification of VAMP2-pHluorin exocytic vesicle fusion events and example maximum projections of inverted time-lapse TIRF images of *Trim9^+/+^* neurons expressing VAMP2-pHluorin treated with netrin-1 or netrin-1 and FAKi. Red circles indicate sites of vesicle fusion events. Montages represent examples of exocytic fusion events in a soma (top) and neurite (bottom). p values from Kruskal-Wallis ANOVA with Bonferroni post-hoc correction to *Trim9^+/+^*control, unless otherwise noted. **C)** Average projection of TIRF time lapse images of GFP-VAMP2 vesicles in *Trim9^+/+^* control and FAKi treated neurons. Quantification of % of the total cell area containing non-motile vesicles. p value from Kruskal-Wallis ANOVA with Tukey post-hoc correction. **D)** Representative transmission electron micrographs of *Trim9^+/+^* control and FAKi treated neurons. Left graph shows density of vesicles within 100 nm of the plasma membrane, p value from Mann-Whitney test. Right graph shows vesicle diameter, p value from t-test.

These data suggested the intriguing possibility that FAK activity functions in the progression from assembled trans-SNARE complexes to lipid bilayer fusion. We hypothesized that during FAKi treatment, these nonproductive SNARE complexes might be tethering non-fusing vesicles to the plasma membrane. To test this, we imaged eGFP-VAMP2 within 100 nm of the basal plasma membrane of *Trim9^+/+^* cortical neurons via TIRF microscopy to visualize vesicles at the basal neuronal surface (**Fig. 6C, Movie 6)**. FAKi treatment was associated with an increased percentage of the basal neuronal membrane area containing non-motile eGFP-VAMP2 in both the presence and absence of netrin-1 (**Fig. 6C**), supporting the hypothesis that FAK activity is required for the progression from assembled SNARE complexes to membrane fusion. To determine if FAKi treatment increased the number of vesicles adjacent to the plasma membrane or altered vesicle size, we performed transmission electron microscopy of cultured neurons. This confirmed an increased density of vesicles within 100 nm of the plasma membrane in FAKi treated neurons, with no change in vesicle diameter (**Fig. 6D**). These data suggest FAK activity is required for the progression from trans-SNARE complex to vesicle fusion. Therefore, by modulating the activation of FAK, TRIM9 controls exocytosis and axon branching.

## Discussion

Here we demonstrate TRIM9-dependent mechanisms that control the localization and function of the netrin receptor DCC and the downstream signaling events and exocytic machinery that promote netrin-dependent axonal branching. Our data suggest that the TRIM9-dependent ubiquitination of DCC that occurs in the absence of netrin-1 precludes activation of FAK and SFKs (**Fig. 7**), important regulators of netrin-dependent axonal morphogenesis. TRIM9 thus maintains the neuron in an immature morphology, yet poised for the rapid responses to netrin-1 required for axon branching. Upon addition of netrin-1, DCC ubiquitination is reduced, and TRIM9 mediates clustering and multimerization of DCC at the plasma membrane and promotes downstream interaction with and activation of FAK and SFKs, followed by elevated SNARE-mediated exocytosis and axon branching. Our studies suggest that this pathway is relevant *in vivo* as well. We identified an unexpected role for FAK activity in the progression from assembled SNAREs to exocytic vesicle fusion at the plasma membrane. Together with previous work demonstrating that TRIM9 constrains the function of the exocytic t-SNARE SNAP25 and the filopodial actin polymerase VASP in the absence of netrin-1 (Li *et al.*, 2001; Mille *et al.*, 2009; Bin *et al.*, 2013; Winkle *et al.*, 2014; Menon and Boyer *et al.*, 2015), these findings reveal an overarching function for TRIM9 in preparing the neuron for spatiotemporally controlled morphological responses to the axon guidance cue netrin-1. By moderating the local signaling necessary for axon branching, TRIM9 plays an integral role in establishing proper morphogenesis *in vitro* and *in vivo*, which is likely critical for behavior, as *Trim9^-/-^* mice exhibit severe deficits in spatial learning and memory (Winkle and Olsen *et al.*, 2016).

**Figure 7.**
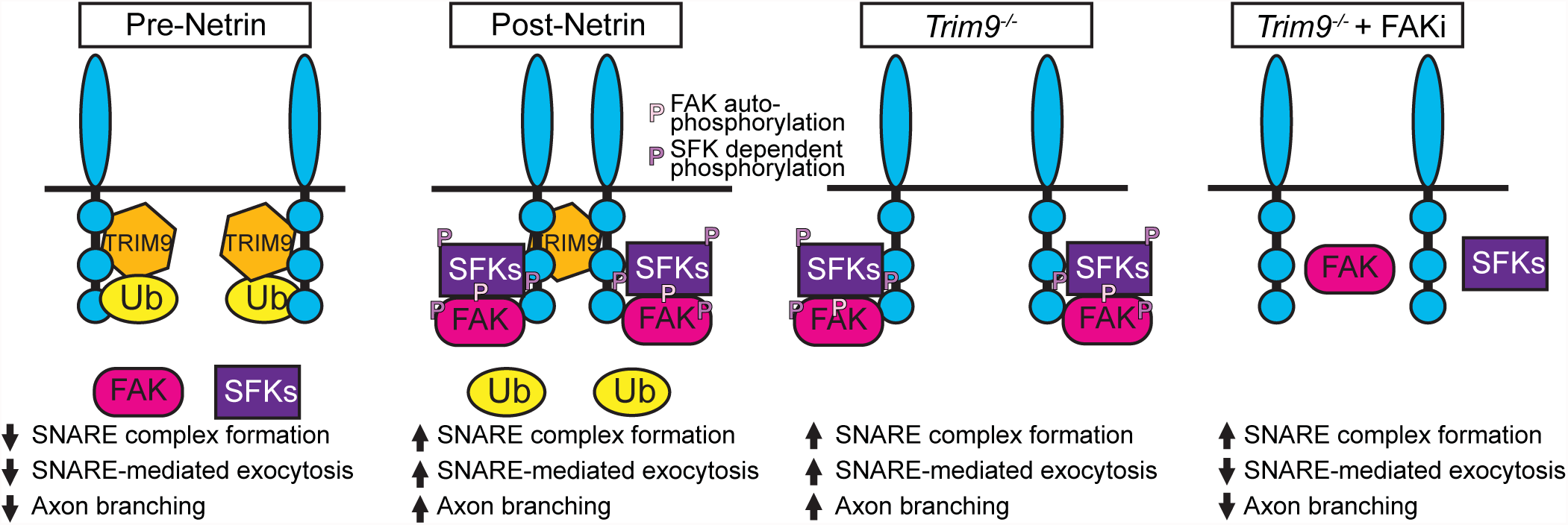
Model of TRIM9 mediated regulation of signaling scaffolds. In the absence of netrin-1, DCC is ubiquitinated in the presence of TRIM9. The addition of ubiquitin inhibits FAK binding and downstream activation of FAK and SFK and constrains vesicle fusion and axon branching. The addition of netrin-1 results in loss of DCC ubiquitination and TRIM9-dependent clustering of DCC. FAK binds to the P3 domain of DCC and is autophosphorylated, leading to interactions with SFK and SFK-dependent phosphorylation events. This creates locally activated signaling complexes that promote vesicle fusion and axon branching. In the absence of TRIM9, the loss of DCC ubiquitination leads to aberrant interaction and activation of FAK and SFK, but a lack of signal complex clustering, leading to aberrant exocytosis and branching. Inhibition of FAK activity blocks subsequent SFK activity, as well as SNARE-mediated exocytosis and branching.

### Non-degradative roles for DCC ubiquitination

Classically, ubiquitination targets a substrate for degradation. Previous studies have demonstrated that DCC ubiquitination and proteasomal degradation of DCC are promoted by addition of netrin (Kim *et al.*, 2005). After the acute netrin-1 treatments used here, ubiquitination of DCC was decreased in cortical neurons, but DCC levels were not. Our findings in both cortical neurons and HEK293 cells support TRIM9-dependent ubiquitination of DCC; however, whether TRIM9 directly ubiquitinates DCC remains unknown. Interestingly, the Siah family of ubiquitin ligases have been implicated in the ubiquitination of both DCC and TRIM9 in heterologous cells (Hu *et al.*, 1997; Fanelli *et al.*, 2004). Although loss of TRIM9 reduced DCC ubiquitination in both neurons and human cells, residual ubiquitination remained, suggesting that DCC may be a substrate of multiple ligases. Alternatively, TRIM9 may regulate the function of another ligase, such as Siah. How these ligases cooperate to regulate DCC remains to be determined.

In neurons, FAK and SFK are activated downstream of DCC binding to netrin-1, analogous to “outside-in” activation of integrin receptors triggered by binding their extracellular ligands. Integrins alternatively can be activated “inside-out” through cytoplasmic binding partners (Shattil *et al.*, 2010; Ye and Petrich, 2011). Our data are consistent with a model in which TRIM9-dependent ubiquitination of the cytoplasmic tail of DCC prevents “inside-out” activation of this signaling pathway by occluding FAK binding to DCC (**Fig. 7)**. Inhibition of the interaction between FAK and DCC has previously been shown to prevent FAK and SFK-dependent phosphorylation of FAK, Fyn, Src and DCC (Li *et al.*, 2004; Liu *et al.*, 2004; Ren *et al.*, 2004). Downstream of DCC pY1418, activation of guanine nucleotide exchange factors (GEFs) that activate Rac1 results in increased actin polymerization (Antoine-Bertrand *et al.*, 2015). This actin polymerization may also participate in the formation of nascent axon branches. This non-degradative function of ubiquitination may represent a new mode of rapid protein and pathway regulation in the cell, that prevents interactions and phosphorylation events.

### Ligase independent function of TRIM9

Here we discovered a ligase-independent function of TRIM9 in the spatial organization of DCC in response to netrin-1. The dramatic and rapid clustering of DCC at the cell periphery that occurred in response to netrin-1 was completely absent in *Trim9^-/-^* neurons. In the *C. elegans* anchor cell, the netrin ortholog *unc-6* stabilizes oscillating clusters of the DCC ortholog *unc-40*, and the TRIM9 ortholog *madd-2* facilitates ligand-independent clustering (Wang *et al.*, 2014), similar to TRIM9ΔRING in mammalian cortical neurons, which suggests evolutionarily conserved functions for TRIM9 in DCC clustering. Structure-function experiments in neurons indicate that the DCC binding and multimerization domains of TRIM9 are necessary for this response, suggesting that the TRIM9 interaction with DCC is responsible for this spatial reorganization. However that interaction alone cannot explain the netrin-dependent clustering as shown by co-immunoprecipitation experiments. Further, the observation that the TRIM9 mutant lacking the ligase domain could induce clustering in the absence of netrin-1 suggests that ligase domain or ligase activity of TRIM9 prevents DCC clustering. This is unlikely through ubiquitination of DCC, as DCC^KR^ did not exhibit aberrant clustering prior to netrin-1 application. Like most E3 ligases, TRIM9 is capable of auto-ubiquitination *in vitro* (Tanji *et al.*, 2010). We speculate that netrin-dependent auto-ubiquitination of TRIM9 could permit TRIM9-dependent clustering of DCC, but this remains to be shown. DCC association with lipid rafts is required for netrin-1 dependent axonal outgrowth (Hérincs *et al.*, 2005). An intriguing hypothesis is that these netrin-dependent DCC clusters represent regulated signaling microdomains possibly within lipid rafts near netrin-1 sources. The formation of these local domains may ensure the spatial and temporal specificity required to form collateral axon branches or may be required for axon turning during development.

### TRIM9 employs a ‘double brake’ on exocytosis and axon branching

Our previous work suggested that TRIM9 sequesters the exocytic t-SNARE SNAP25, preventing SNARE complex assembly. Upon addition of netrin-1, TRIM9 releases SNAP25, which permits SNARE complex formation, vesicle fusion, plasma membrane expansion, and axon branching (Winkle *et al.*, 2014). Here we identify a second, FAK-dependent mechanism that occurs after SNARE complex assembly yet prior to vesicle fusion, by which TRIM9 restrains membrane expansion. Our findings that FAK activity is necessary for vesicle fusion but not for SNARE complex assembly suggest that FAK-dependent phosphorylation of an unknown substrate regulates the transition from assembled SNARE complexes to lipid bilayer fusion. Whether this is by promoting progression from trans-SNARE to cis-SNARE complex, through promoting fusion pore opening, or by another mechanism is not known. Although no pY residues have been identified in the classical neuronal SNARE components, pY residues have been identified on a number of synaptic vesicle associated proteins, such as the Ca^++^-sensor synaptotagmin (Ballif *et al.*, 2008; Wiśniewski *et al.*, 2010; Trinidad *et al.*, 2012). During synaptic vesicle release, calcium-bound synaptotagmin displaces complexin from the SNARE complex, allowing progression to the final stages of vesicle fusion (Giraudo *et al.*, 2009). Whether a phosphorylation-dependent switch in synaptotagmin function or another exocytic component could be involved in developmental exocytosis in neurons is an exciting possibility. An alternative, intriguing possibility is that FAK-dependent modulation of the small Rho GTPase Cdc42, which functions in a late stage of exocytosis in yeast (Adamo *et al.*, 2001; Myers *et al.*, 2012), could be regulating vesicle fusion in neurons.

Taken together, our data indicate that FAK activity is necessary for netrin-dependent increases in exocytosis and axon branching, and that the elevated FAK activity that occurs in the absence of *Trim9* is required for the elevated exocytosis and axon branching in these neurons. Intriguingly, the relevance of this pathway *in vivo* is supported by our finding that expression of a noncatalytic version of FAK reduces the exuberant axon branching in the corpus callosum of *Trim9^-/-^* mice. Our previous studies indicate that TRIM9 is also required for netrin-dependent axon guidance (Menon and Boyer *et al.*, 2015). Application of the repulsive cue Sema3a decreases VAMP2-mediated exocytosis and Sema3a receptor trafficking, which regulates repulsive growth cone turning (Zylbersztejn *et al.*, 2012). Analogously, we speculate that TRIM9 and FAK may regulate axon guidance via the modulation of VAMP2-mediated exocytosis.

## Methods

### Animals

All mouse lines were on a C57BL/6J background and bred at UNC with approval from the Institutional Animal Care and Use Committee. Timed pregnant females were obtained by placing male and female mice together overnight; the following day was designated as embryonic day (E) 0.5 if the female had a vaginal plug. *Trim9^-/-^* and Thy1-GFP (Line M) mice were as previously described, (Feng *et al.*, 2000; Winkle *et al.*, 2014). We crossed the *Trim9^-/-^/*Thy1GFP mice, with Nex-Cre mice (Goebbels *et al.*, 2006) and FRNK^loxp^ mice (DiMichele *et al.*, 2009) (generous gift from Dr. Joan Taylor (University of North Carolina at Chapel Hill).

### Antibodies, Reagents and Plasmids

Antibodies used in this study include mouse anti-Myc (9E10), rabbit anti-FLAG (Sigma), mouse anti-HA (Millipore), rabbit anti-GFP (Invitrogen), mouse anti-GFP (Neuormab), rabbit anti-βIII tubulin (Covance), mouse anti-βIII tubulin (Covance), mouse anti-GAPDH (SCBT), mouse anti-Ubiquitin (SCBT), mouse anti-Syntaxin 1A (SCBT), rabbit anti-VAMP2 (Cell Signaling), goat anti-SNAP25 (SCBT), rabbit anti-FAK-pY397 (Invitrogen), rabbit anti-FAK-pY861 (Novex), mouse anti-FAK (Invitrogen), goat anti-DCC (A-20, SCBT), and rabbit anti-DCC pY1418 (generously provided by Dr. Nathalie LaMarche Vane, McGill University). Anti-Myc (9E10) conjugated agarose beads (SCBT), DCC-A20 conjugated agarose beads (SCBT) or HA antibody conjugated sepharose beads (Cell Signaling) were used for immunoprecipitations. AlexaFluor 488, AlexaFluor 568, or AlexaFluor 647 labeled secondary antibodies and phalloidin (Invitrogen) were used for immunocytochemistry. Far-red labeled secondary antibodies (Licor) were used for western blots.

Fluorescent and epitope tagged TRIM9 mammalian expression plasmids have been previously described (Winkle *et al.*, 2014). To generate pHluorin-DCC, pHluorin was subcloned into mCherry-DCC (Winkle *et al.*, 2014). High-confidence ubiquitination sites were predicted using UbPred (Radivojac *et al.*, 2010), and the DCC^KR^ mutant was generated via GENEBlocks (Life Technologies). pHluorin-VAMP2, eGFP-VAMP2 and eGFP-VAMP7 were described (Miesenböck *et al.*, 1998; Gupton and Gertler, 2010). HA-DCC and Myc-DCC were generous gifts from Dr. Tim Kennedy (Mcgill University). Myc-FAK was a generous gift from Dr. Joan Taylor (UNC-Chapel Hill). The rat FRNK construct was a kind gift of J.T. Parsons (UVA, Charlottesville, VA). Recombinant chicken netrin-1 was concentrated from supernatant of HEK293 cells (Serafini *et al.*, 1994; Lebrand *et al.*, 2004). The FAK inhibitor 14 (Tocris), Bortezomib (SCBT) and MG132 (American Peptide Co) were used at the indicated concentrations.

### Cell Culture and Transfection

E15.5 dissociated cortical neuronal cultures were prepared as described (Viesselmann *et al.*, 2011). Briefly, cortices were micro-dissected and neurons were dissociated with trypsin and plated on Poly-D-lysine (Sigma)-coated coverslips, glass bottom movie dishes (Matek) or tissue culture treated plastic in Neurobasal media (Invitrogen) supplemented with B27 (Invitrogen). For all plasmid transfections, dissociated neurons were resuspended in mouse neuron Lonza Nucleofector solution and electroporated in accordance with the manufacturer protocol prior to plating. *TRIM9^+/+^* and CRISPR-generated, *TRIM9^-/-^* HEK293 cells were described previously (Menon and Boyer *et al.*, 2015). HEK293 cells were maintained in DMEM with glutamine (Invitrogen) supplemented with 10% FBS (Hyclone). HEK293 cells were transfected using Lipofectamine 2000 (Invitrogen) as per manufacturer protocol.

### Immunoblotting and Immunoprecipitation

For all immunoblotting and quantification SDSPAGE and immunoblot analysis were performed using standard procedures. All lysate concentrations were calculated via spectrophotometry utilizing Bradford concentration reagent (Biorad). Samples were diluted in 5X sample buffer, resolved via SDS-PAGE, transferred to nitrocellulose, and the resulting western blots were probed with primary and corresponding secondary antibodies as indicated. Signal was detected with Odyssey Imager (Licor) and either analyzed via Licor Image studio lite or ImageJ. All values were normalized to loading control and to wildtype control condition, unless otherwise noted.

Co-immunoprecipitations were performed using mouse IgG-conjugated A/G beads (SCBT) to pre-clear lysates for 1 h at 4°C with agitation. For DCC multimerization studies, cells were co-transfected with HA-DCC and pHluorin- (GFP) DCC, treated and lysed in lysis buffer (50 mM Tris pH7.5, 200 mM NaCl, 2 mM MgCl_2_, 10% glycerol, 1% NP-40, supplemented with protease and phosphatase inhibitors). HA-DCC and interacting proteins were immunoprecipitated from cell lysate overnight at 4°C. For FAK co-immunoprecipitation with pHluorin-DCC, cells were transfected with Myc-FAK and either pHluorin-DCC or pHluorin-DCC^KR^, treated and lysed in lysis buffer. Lysates were incubated with DCC antibody overnight at 4°C. For DCC co-immunoprecipitation with Myc-TRIM9, cells were transfected with HA-DCC and Myc-TRIM9, treated, and lysed in lysis buffer. Lysates were incubated with Myc antibody overnight at 4°C. Beads were washed three times with lysis buffer and bound proteins were prepared in sample buffer, resolved by SDS-PAGE and analyzed by immunoblotting.

To measure ubiquitination, HEK293 cells transfected with HA-DCC, pHluorin-DCC, or pHluorin-DCC^KR^ and FLAG-Ub or cortical neurons at 48 h after plating were treated with 10 μM MG132 for 4 hr +/- 600 ng/ml netrin-1for 40 min. Cells were lysed in ubiquitin immunoprecipitation buffer (20 mM Tris-Cl, 250 mM NaCl, 3 mM EDTA, 3 mM EGTA, 0.5% NP-40, 1% SDS, 2 mM DTT, 5 mM NEM (N-ethylmaleimide), 3 mM iodoacetamide, protease and phosphatase inhibitors pH=7.3-7.4). For 5-6 million cells 270 μl of ubiquitin immunoprecipitation buffer was added and incubated on ice for 10 min. Cells were removed from the dish and transferred into tubes. 30 μl of 1X PBS was added and gently vortexed. Samples were boiled immediately for 20 min until clear, then centrifuged at 14,000 rpm for 10 min. The boiled samples were diluted using IP buffer without SDS to reduce SDS concentration to 0.1%. DCC from HEK293 cells was immunoprecipitated using the appropriate tags. A similar protocol was used for untransfected cultured neurons, using antibodies against endogenous DCC and ubiquitin. Immunoprecipitations were performed under conditions in which only covalent interactions were preserved with anti-GFP (mouse)/ anti-HA (mouse) antibody or anti-DCC (goat) antibody coupled to Protein A/G agarose beads.

To assess proteasomal degradation of DCC, *Trim9^+/+^* and *Trim9^-/-^* E15.5 cortical neurons were treated with 100 nM bortezomib for 4 h prior to 40 min of 500 ng/ml netrin-1 stimulation and lysis. To assess the FAK pY397 and pY861, in response to netrin-1 stimulation, 250 ng/ml of netrin-1 was bath applied at the indicated times prior to lysis, or left untreated for control. To assess total DCC and DCC pY1418 protein levels, 500 ng/ml of netrin-1 was bath applied between at 72 h to dissociated *Trim9^+/+^* and *Trim9^-/-^* E15.5 cortical neurons for 30 min prior to lysis or left untreated for control. Neurons were lysed in a lysis buffer and stabilized in SDS sample buffer and processed and quantified as described above. To assess SNARE complexes, assays were performed as extensively outlined (Winkle *et al.*, 2014; 2016a). In brief, neuronal cell lysates from untreated, netrin-1 treated (250 ng/mL) or FAKi treated (5 μM) for 1 h and were stabilized in sample buffer and incubated at 37°C for complexes or 100°C for monomers. Resulting samples were processed and quantified as described above.

### Live Cell Imaging

Live cell confocal z stacks of mCherry-DCC were acquired at RT with a spinning disk confocal (Yokogawa CSU-X1) on an inverted Olympus IX81 zero-drift microscope with Andor iQ acquisition software (Andor Revolution XD system), an electron-multiplying charge-coupled device (iXon), a 50 mW 561 nm laser, band pass emission filters (Semrock brightline) for 607–36 nm (TR-F607-036), and a 60x PlanSApo 1.2 W objective. Live cell TIRF imaging was performed on an inverted microscope (IX81-ZDC2) with MetaMorph acquisition software, an electron-multiplying charge-coupled device (iXon), and an imaging chamber (Stage Top Incubator INUG2-FSX; Tokai Hit), which maintained 37°C and 5% CO_2_.

To measure DCC spatial clustering, neurons transfected with mCherry-DCC, pHluorin-DCC or mCherry-DCC and GFP-TRIM9 variants were imaged every 10 s for 8 min before and after stimulation with 500 ng/ml netrin-1, with a 100x 1.49 NA TIRF objective and a solid-state 491 nm and 561 nm laser illumination at 120 nm penetration depth at 48 h *in vitro*. Resulting image stacks were aligned using the ImageJ plugin (TurboReg). For analysis of DCC clustering, cell masks were automatically generated using a cell segmentation algorithm in MATLAB. The average of the pre-treated image stacks were subtracted from each image followed by a sharpening mask, a Gaussian blur, and finally a threshold was applied using Otsu’s method (Otsu, 1979) to create the cell mask. This cell mask was sub-divided into an edge mask, defined as the outer pixels nearest the cell perimeter (0.64 μm in width) and the inner mask, defined as the remaining pixels that did not belong to the edge mask. DCC fluorescence intensity was corrected for photobleaching using the ImageJ CorrectBleach plugin. Image stacks were temporally aligned to the “treatment frame” at which time the cells were stimulated with netrin. The average pixel intensity for each region of interest (whole cell, perimeter, and cell center) was calculated for each time point and normalized to the whole cell fluorescence. The temporal changes in average DCC fluorescence intensity for each ROI were fit using local polynomial regression fitting (LOESS). All LOESS curves were second order polynomials fit with α = 0.75. For comparison of the brightest DCC-containing pixels (assumed to represent the most clustered DCC molecules) the pixel intensities of the pre-treatment image stack and posttreatment stack were averaged. A threshold was applied to capture the top 10% of brightest pixels in the cell mask. The percentage of these pixels that were in the cell edge mask was compared across each condition.

For exocytic vesicle fusion assays, neurons expressing VAMP2-pHluorin were imaged at 48 h after plating with a 100x1.49 NA TIRF objective and a solid-state 491 nm laser illumination at 100 nm penetration depth. Images were acquired every 0.5 s for 5 min. The frequency of exocytic events normalized per cell area and time is reported for the entire cell area. 500 ng/ml netrin-1 +/- 1 μM FAKi was added to the dish 1 h before imaging of netrin-1-stimulated neurons. For imaging eGFP-VAMP2 containing vesicle motility, *Trim9^+/+^* neurons expressing eGFP-VAMP2 and cytoplasmic mCherry were imaged at 100x magnification every 2 s for 5 min at 48 h after plating at 110 nm TIRF penetration depth. FAKi treatment (5 μM) occurred 30 min prior to imaging. Images were analyzed using ImageJ (Schindelin *et al.*, 2012); a threshold of the mCherry images was used to measure the total area of the footprint of the cell, and an average projection of the eGFP-VAMP2 time-lapse was used to identify non-motile vesicles. The percentage of the neuronal footprint area containing non-motile vesicles is reported.

### Axon Branching Assays

For *in vitro* axon branching assays, 250 ng/ml netrin-1 was bath-applied after 48 h in culture, and FAK inhibited conditions were treated with 1 μM FAK inhibitor. Cells were fixed in 4% PFA for 20 min at 72 h in culture, permeabilized for 10 min in 0.1% Triton X-100, blocked for 30 min in 10% BSA, and incubated with indicated primary antibodies for 1 h at RT. Following three washes, neurons were incubated in spectrally distinct, species appropriate, fluorescent secondary antibodies for 1 h at RT. Following three washes, cells were mounted in a TRIS/glycerol/n-propyl-gallate-based mounting media for imaging. Widefield epifluorescence images of pyramidal-shaped neurons were collected at RT on an inverted microscope with 40x 1.4 NA Plan Apochromat objective lens (Olympus), MetaMorph acquisition software, and an electron-multiplying charge-coupled device (iXon). Axon branches were defined as extensions off the primary axon that were ≥ 20 μm in length.

For analysis of axon branching *in vivo,* 200 μm coronal sections of 3-week-old Thy1-GFP/*Trim9^+/+^,* Thy1-GFP/*Trim9^+/+^*/Nex-CRE/FRNK^LoxSTOPLox^, Thy1-GFP/*Trim9^-/-^,* Thy1-GFP/*Trim9^-/-^*/Nex-CRE/FRNK^LoxSTOPLox^ littermates mounted in DPX mountant (Fisher) were imaged on a confocal inverted microscope (Fluoview FV1200, Olympus) equipped with a 20x, 0.75 NA Plan Apochromat objective lens with a 488 nm argon laser. Multi-area z stacks were stitched, ROIs containing potential axon branch points were identified in maximum projections. Candidate branches were confirmed by inspection of stitched z stacks by the intersection and coalescence of two resolved axons. Candidate branches were rejected if axons were observed to intersect but not coalesce or if one axon left the focal plane without having reached the potential branch point.

### Transmission electron microscopy

For EM, neurons were plated on ACLAR Film in 35 mm dishes and prepared as described (Heckman, 2008). Briefly, after 48 hours *in vitro*, neurons were left untreated or a 30 minute treatment with 5 μM FAKi. Subsequently they were rinsed with 1X PBS pH 7.5, fixed for 20 minutes in 0.1 M NaCL, 0.05 M cacodylate pH7.5 containing 2.5% w/v glutaraldehyde, washed two times PBS, and postfixed in a solution containing 1% w/v osmium tetroxide and 1.5% w/v potassium ferricyanide. After dehydration in graded hexylene glycol, cells were embedded in plastic (Spurr’s resin with ERL-4221). Thin sections (70 nm) were cut and collected on nickel mesh grids, and contrasted with 4% w/v uranyl acetate followed by 0.4% w/v lead citrate. Sections were examined under a transmission electron microscope (Philips Tecnai 12). Circular structures with a membrane were considered vesicles. All vesicle diameters were measured. Vesicles within 100 nm of the plasma membrane were reported. All analysis was performed blinded.

### Statistics

At least three independent experiments were performed for each assay. Data distribution normality was determined using the Shapiro-Wilkes test. Normally distributed data were compared by unpaired *t-*test for two independent samples. For more than two samples with normal distribution, statistical comparisons were made by Analysis of Variance (ANOVA) with Bonferroni (3-4 comparisons) or Tukey (>4 comparisons) post-hoc correction. For non-normally distributed data, the Mann-Whitney test was used to compare two samples. For more than two samples, Kruskal-Wallis non-parametric ANOVA with the post-hoc corrections described above was used. For analysis of western blots, Kruskal-Wallis non-parametric ANOVA with LSD correction were made. Data are presented as box-and-whisker plots indicating the 25th percentile (bottom boundary), median (middle line), 75th percentile (top boundary), nearest observations within 1.5 times the interquartile range (whiskers). Outliers are defined as data points outside 1.5 times the interquartile range. Red lines indicate arithmetic means, individual blue data points are included when experimental observations per condition are less than 10, such as western blots. Outliers are included in statistical analyses.

### Online Supplemental Material

Fig S1 shows netrin dependent clustering of DCC occurs on the apical membrane. Movie 1 shows shows netrin-dependent clustering of mCherry-DCC is present in *Trim9^+/+^* neuron and reduced in *Trim9^-/-^* neuron. Movie 2 shows expression of TRIM9 in *Trim9^-/-^ neuron* rescues netrin-dependent clustering of mCherry-DCC, that TRIM9ΔRING causes netrin-independent clustering of mCherry-DCC, and that TRIM9ΔSPRY and TRIMΔCC fail to rescue netrin-dependent clustering of mCherry-DCC. Movie 3 shows netrin-dependent clustering of pHluorin-DCC^KR^. Movie 4 shows FRNK-dependent rescue of aberrant axon branching in the corpus callosum of Thy1-GFP/*Trim9^-/-^* mice. Movie 5 shows VAMP2-pHluorin based exocytic events in netrin-1 treated and netrin-1/FAKi treated neurons. Movie 6 shows that VAMP2-containing vesicle motility is reduced by FAKi treatment.

## Acknowledgements

We thank Patrick Brennwald for thought critique of the manuscript. We thank Gerald Gordon, Robert Currin, Jennifer Marks and Lora Hartman and the UNC Olympus Center for their expertise and generosity. We thank Carey Hanlin for technical assistance. This work was supported by the NIH: GM108970 (S.L.G.), MH10965301 (S.L.G.), F31 NS087837 (C.C.W.), and American Heart Association: 14POST20450085 (S.M.)

